# The crystal structure of the TonB-dependent transporter YncD reveals a positively charged substrate binding site

**DOI:** 10.1101/2020.01.29.925891

**Authors:** Rhys Grinter, Trevor Lithgow

**Affiliations:** School of Biological Sciences, Monash University, Clayton 3800, Victoria, Australia; Infection and Immunity Program, Biomedicine Discovery Institute and Department of Microbiology, Monash University, Clayton 3800, Victoria, Australia

**Keywords:** YncD, Crystallography, Membrane Transport, Gram-Negative Bacteria, TonB-Dependent Transporter

## Abstract

The outer membrane of Gram-negative bacteria is highly impermeable to hydrophilic molecules larger than 600 Da, protecting these bacteria from toxins present in the environment. In order to transport nutrients across this impermeable membrane, Gram-negative bacteria utilise a diverse family of outer-membrane proteins called TonB-dependent transporters. The majority of this family transport iron-containing substrates. However, it is becoming increasingly clear that TonB-dependent transporters target chemically diverse substrates. In this work, we investigate the structure and phylogenetic distribution of the TonB-dependent transporter YncD. We show that while YncD is present in some enteropathogens including *E. coli* and *Salmonella* spp., it is also widespread in Gamma and Betaproteobacteria of environmental origin. We determine the structure of YncD, showing that despite a distant evolutionary relationship, it shares structural features with the ferric-citrate transporter FecA, including a compact positively-charged substrate-binding site. Despite these shared features, we show that YncD does not contribute to the growth of *E. coli* in pure culture under-iron limiting conditions or with ferric-citrate as an iron source. Previous studies on transcriptional regulation in *E. coli* show that YncD is not induced under iron-limiting conditions and is unresponsive to the Ferric uptake regulator (Fur). These observations combined with the data we present, suggest that YncD is not responsible for the transport of an iron-containing substrate.

## Introduction

The outer membrane of Gram-negative bacteria represents a formidable permeability barrier to hydrophilic molecules larger than ~600 Da (1). Hydrophilic molecules smaller than this diffusion limit cross the outer membrane through proteins called porins, which can be either promiscuous or substrate-specific (2). To import molecules that are too large for diffusion through outer-membrane porins, bacteria employ a diverse family of membrane proteins termed TonB-dependent transporters (TBDTs) (3). These transporters in the outer membrane are so named because they engage in active transport, the energy for which is provided by the TonB-ExbBD complex in the inner membrane, which spans the periplasm to interact with TBDTs (4,5). TBDTs are specific for a particular substrate, which they bind with high affinity in an extracellular binding pocket (6,7). This high-affinity binding allows TBDTs to capture molecules that are of low abundance in the environment, enabling bacteria to scavenge scarce nutrients (6). One such scarce nutrient is iron, which is the substrate for the majority of characterised TBDTs (3). Most iron-transporting TBDTs bind Fe^3+^ ions in complex with siderophores, organic molecules that bind metal ions with high affinity and are secreted by the bacteria or by competitors from the microbial community (8). In addition to importing Fe-siderophore complexes, some TBDTs harvest iron directly from proteins (9,10). These protein-binding TBDTs often target host proteins and are employed by pathogens to obtain iron during infections (11,12).

While many of the most intensively studied TBDTs import iron-containing substrates, it is becoming increasingly clear that members of the TBDT protein family are responsible for the import of chemically diverse substrates (13). Non-iron TBDT substrates range in size from free metal ions, like Cu^2+^ and Zn^2+^, to cobalamin, polysaccharides, polypeptides, and small globular proteins (9,14–22). As functionally characterised TBDTs represent a small segment of this protein family, the extent of the substrate diversity of this family remains to be discovered.

The model bacterium *Escherichia coli* K12 possesses 9 TBDTs which target different substrates for import (7). The TBDTs FepA, CirA and Fiu transport Fe^3+^ containing catecholate siderophores (7,23,24), FhuA and FhuE transport Fe^3+^ containing hydroxamate siderophores of fungal origin (6,25), FecA transports iron citrate (26), while BtuB targets cobalamin (16). Recent research suggests that the TBDT YddB from *E. coli* K12 functions in the import of a small iron-containing protein, although the exact nature of this substrate remains to be identified (27). The final TBDT possessed by *E. coli* K12 is YncD, and the substrate for this transporter remains unknown. The homolog of YncD in *Salmonella enterica* ssp. *enterica* serovar Typhi has been shown to be important for virulence in a mouse model of infection (28). As such, characterisation of YncD is important for understanding infections caused by *Salmonella* and other enterobacterial pathogens that possess it.

In this study, we investigated the structure and phylogenetic distribution of YncD. Using a comparative genomics approach, we show that YncD is present in members of Enterobacteriaceae that form commensal or pathogenic associations with the human intestine, and is more broadly present in environmental Proteobacterial isolates, including species which are opportunistic pathogens of humans. We determined the crystal structure of YncD demonstrating that it possesses a compact positively-charged extracellular substrate-binding site that is characteristic of TBDTs that import negatively charged small-molecule substrates. Finally, using an *E. coli* strain deficient in all iron transporting TBDTs we show that YncD does not play a role in iron acquisition under laboratory conditions, providing further evidence that this transporter does not transport an iron-containing substrate.

## Results

### The TonB-dependent transporter YncD is widespread in environmental bacteria

To determine the evolutionary relationship between YncD and other functionally characterised TBDTs, we performed clustering analysis of these sequences based on pairwise sequence similarity scores, using the program CLANS (29). This analysis reveals that YncD is distantly related to other TBDTS of known function and does not form a cluster with any of these sequences (Figure 1A). This analysis shows that while YncD is most closely related to the TBDTs FecA and BtuB, this reflects only 21% and 18% sequence identity, respectively.

**Figure 1.**
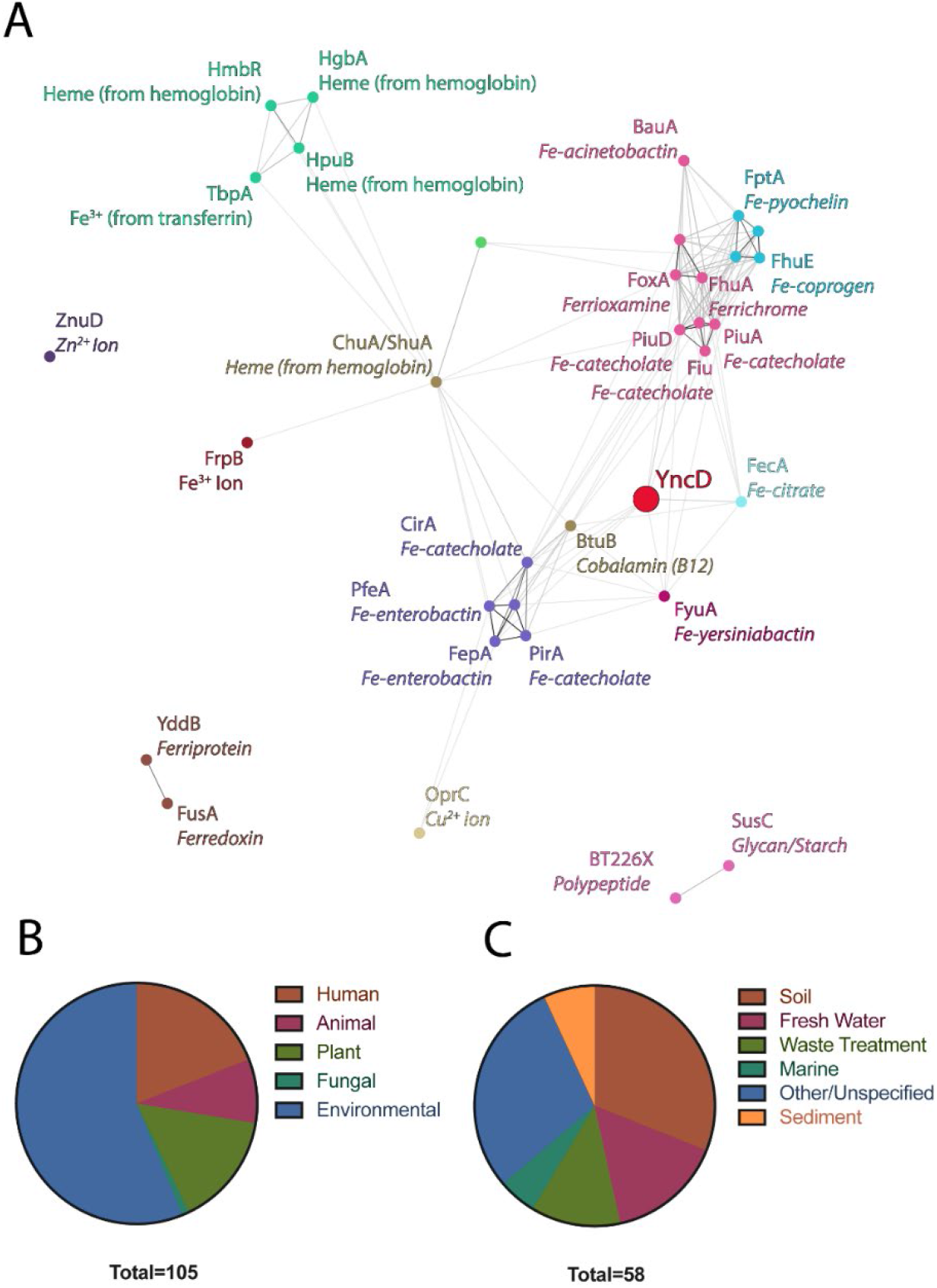
The relationship between YncD and characterised TBDTs and the origin of bacteria that possess YncD. (A) Sequence clustering analysis (E-value cutoff = 1×10^−10^) of YncD with TBDTs of known structure and/or function. Sequences are colored-coded according to the sequence cluster to which they belong. The circle representing the sequence of YncD is enlarged and colored red, showing that YncD does not cluster with other sequences. (B) The environment or host of isolation of a subset of the YncD containing bacteria summarised in Table S1. (C) The environment type of YncD containing bacterial isolates from environmental sources.

To gain insight into the distribution of YncD in Gram-negative bacteria, we interrogated the Uniprot reference proteomes for YncD homologues with the HMMER search algorithm (30,31). An initial sequence similarly E-value cut-off of 1×10^−50^ was used to capture all YncD sequences present in the database. To remove false-positive non-YncD sequences from this dataset, we subsequently clustered the sequences using CLANS (29) with an E-value cut-off of 1×10^−130^. The largest sequence cluster had 359 distinct sequences including YncD from *E. coli* and was extracted for curation and analysis (Table S1, Figure S1A). Partial sequences were removed from this dataset, yielding 331 sequences. These were reclustered with an E-value cut-off of 0, revealing several distinct sub-sets within the YncD protein sub-family (Figure S1B). A multiple sequence alignment of these diverse YncD sequences was then performed revealing that they share amino acid sequence identities ranging between 33 and 99 % (Table S2, Data S1), and highlighted a number of conserved regions. Most notably the region of the corresponding in amino acids 79-150 of YncD from *E. coli* is highly conserved (Data S1). This region encompasses the extracellular loops of the N-terminal plug domain of the transporter, known to mediate substrate binding, suggesting that the YncD homologues identified target a chemically analogous substrate.

To explore potential niche-specific functions associated with YncD, we interrogated genome metadata from those bacteria identified in the YncD HMMER search. The environment of isolation of 105 YncD-containing species was determined, with all bacteria belonging to either the gamma- or beta-proteobacteria (Table S1). While some of these bacterial species were isolated from a human host (largely Enterobacteriaceae associated with the gut or Pseudomonadales associated with respiratory infection), the majority of YncD-containing bacteria were environmental isolates (Figure 1B). The environmental isolates originated from soil, sediment or water samples (Figure 1C). The widespread distribution of YncD in environmental isolates, and its absence from most specialist bacterial pathogens, would be most consistent with YncD playing a role in the acquisition of a molecule produced by the microbial community.

### The crystal structure of YncD from *E. coli* reveals a positively charged substrate-binding site

To gain insight into the evolutionary relationship between YncD and other TBDTs and to provide insight into the nature of the YncD substrate, we solved the crystal structure of YncD by X-ray crystallography. Crystals of YncD from *E. coli* were prepared and diffraction was collected to 3.2 Å (Table 1). Molecular replacement using the closest YncD homologue FecA (PDB ID: 1KMO, 1KMP) proved unsuccessful (26). To obtain experimental phases, YncD crystals were soaked with potassium tetranitroplatinate and anomalous diffraction was collected to 3.5 Å (Table 1). Phases were obtained for this dataset using single-wavelength anomalous dispersion (SAD) phasing, with an anomalous substructure of 2 high occupancy and 1 low occupancy platinum atoms located (Figure S3). An initial model of YncD was build using maps from this dataset and utilised to obtain phases for the higher resolution dataset by molecular replacement. The structure of YncD was then built and refined using the native 3.2 Å data (Table 1).

**Table 1.**
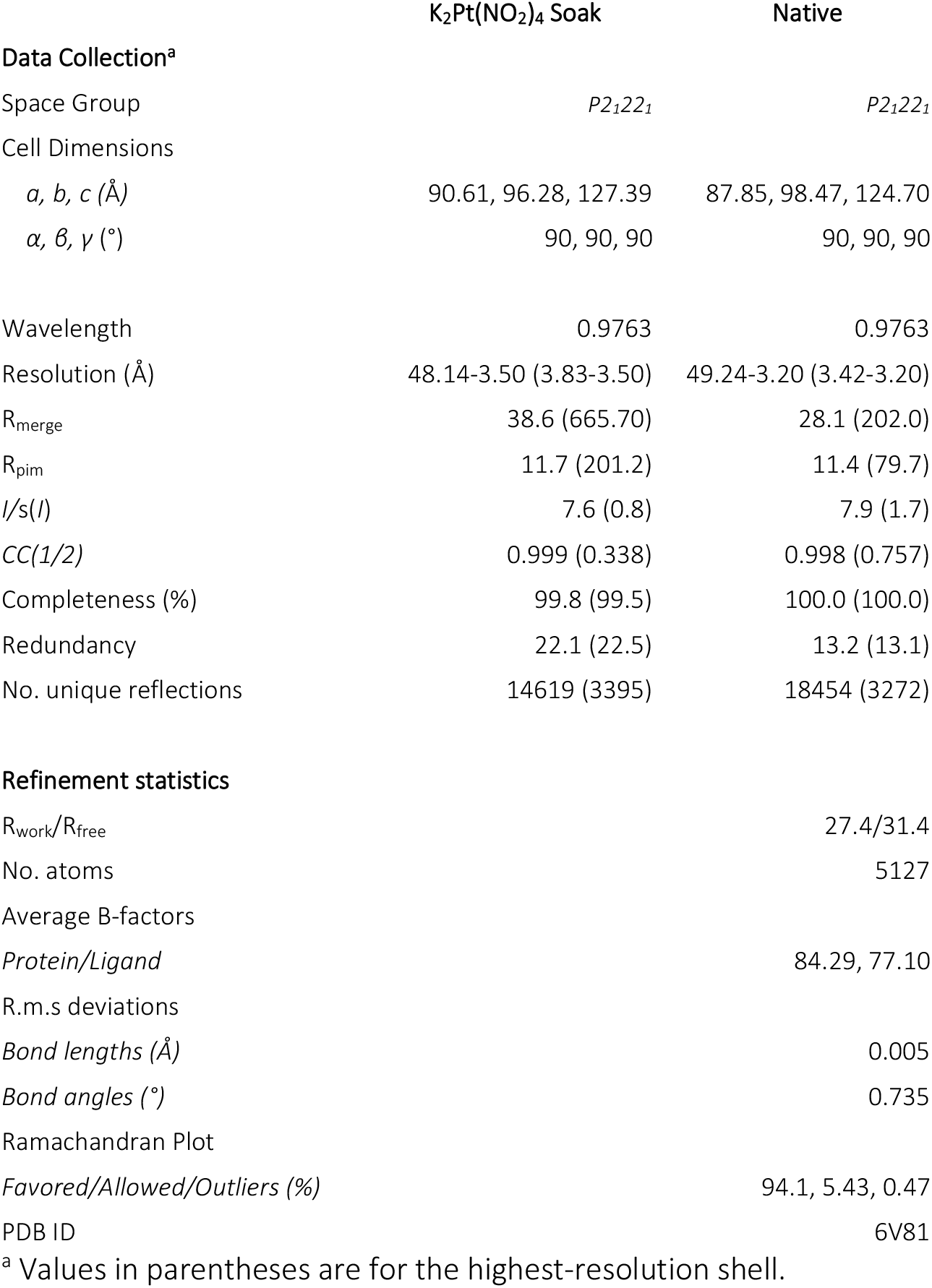
YncD crystallography data collection and refinement statistics.

As expected, based on other members of the TBDT family, structurally YncD consists of at 22 stranded transmembrane β-barrel, which is occluded by a globular N-terminal plug domain (Figure 2A, B). The Dali server was utilised to search the protein databank (PDB) for structural homologues to YncD (32), revealing that consistent with our sequence analysis the closest structural homologue to YncD is FecA (Table S3). Despite the low sequence identity of 21 % between YncD and FecA, these proteins share a backbone atom root mean square deviation (RMSD) of 2.2 Å. Consistent with this, the extracellular loop length, secondary and tertiary structure is relatively well conserved between the two proteins, compared to the more distantly related TBDT FhuE (Figure 2C, Figure S2).

**Figure 2.**
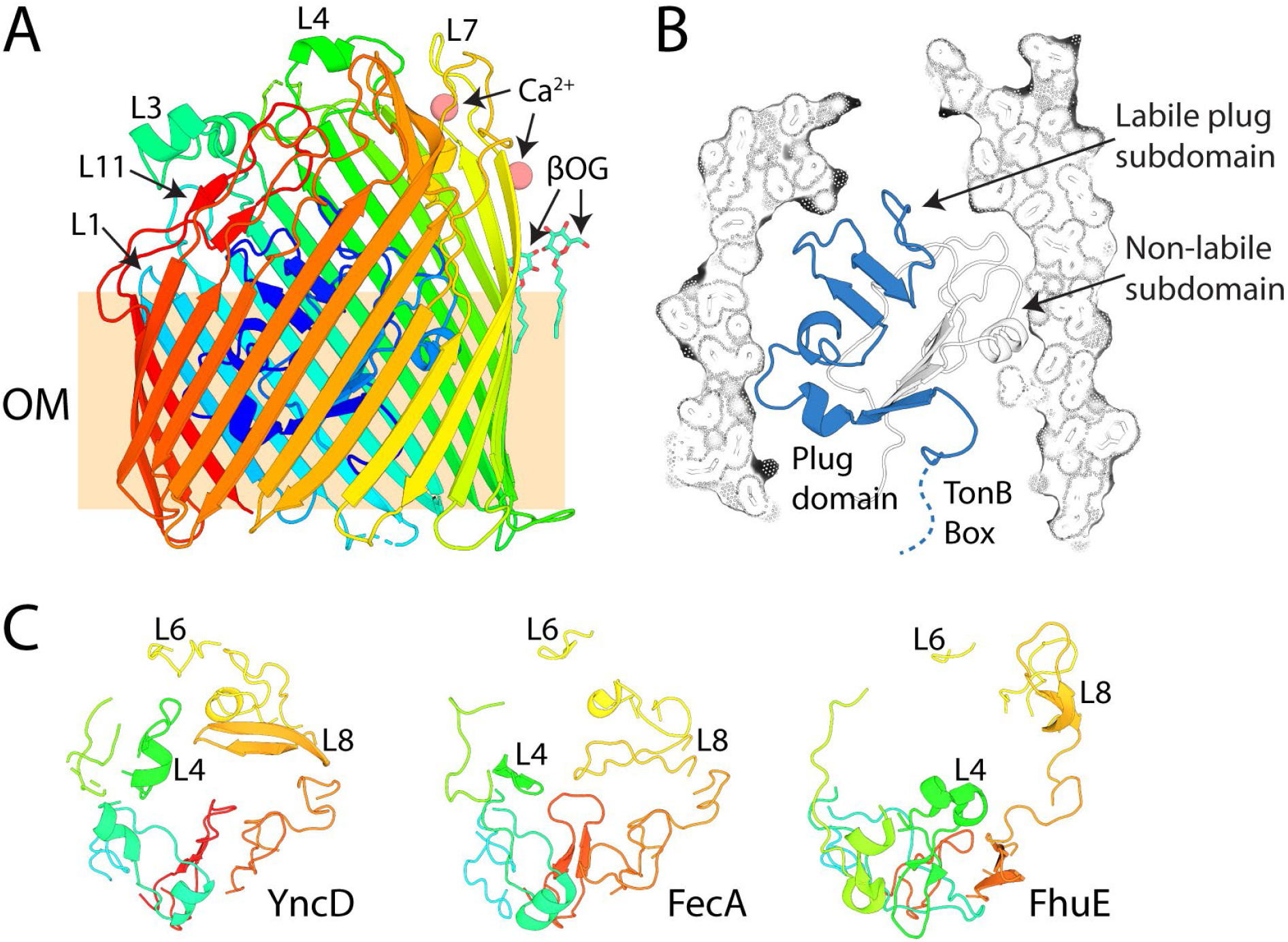
The crystal structure of YncD reveals structural homology to the distantly-related transporter FecA. (A) The crystal structure of YncD shown as a cartoon representation with rainbow coloring from the N-terminus (blue) to the C-terminus (red). Selected loops are labelled for reference i.e. L3 = Loop 3. The outer membrane-embedded region is boxed and with the label ‘OM’. Ca2^+^ ions and βOG detergent molecules co-crystallised with YncD are shown and sphere and stick representations respectively. (B) The crystal structure of YncD with the barrel domain shown as a cutaway surface and stick representation and the plug-domain shown as a cartoon representation. The labile section of the plug domain displaced by TonB is shown in blue, including the TonB box, which is disordered in the structure. (C) The extracellular loops of YncD, FecA (PDB ID = 1KMP) and FhuE (PDB ID = 6E4V) shown as a cartoon representation, with rainbow coloring as in panel A.

Previous analysis of the structure of diverse TBDTs in complex with their substrates revealed the location of a conserved substrate-binding site in these transporters (7). Based on this analysis we identified a compact substrate-binding site formed by the extracellular loops of YncD in our structure (Figure 3A). Analysis of sequence conservation among YncD homologues using the program Consurf (33) and our alignment of YncD sequences revealed that the residues that constitute this binding pocket are well conserved in the YncD homologues, suggesting they play an important role in transporter function (Figure 1B, C, Data S1). These conserved residues include a number of arginine, glutamine and lysine residues that give the substrate-binding site of YncD a highly positive charged, based on a calculated electrostatic surface (Figure 3B-D). Electrostatic analysis of the structure of FecA shows that this distant YncD homologue also possesses a highly positively charged substrate-binding site (Figure 3C). In FecA, several arginine and glutamine residues, which give the substrate-binding site its positive character, are responsible for coordinating the carboxylic acid groups of the Fe-citrate complex, and thus are important for substrate binding by FecA (Figure 3C) (26). Superimposition of the FecA-Fe-Citrate complex with YncD places Fe-citrate ligand centrally in the positively charged substrate binding site of YncD (Figure 3C). The similarities between the character of the substrate bindings sites of YncD and FecA suggest that despite their differences they may target chemically similar substrates.

**Figure 3:**
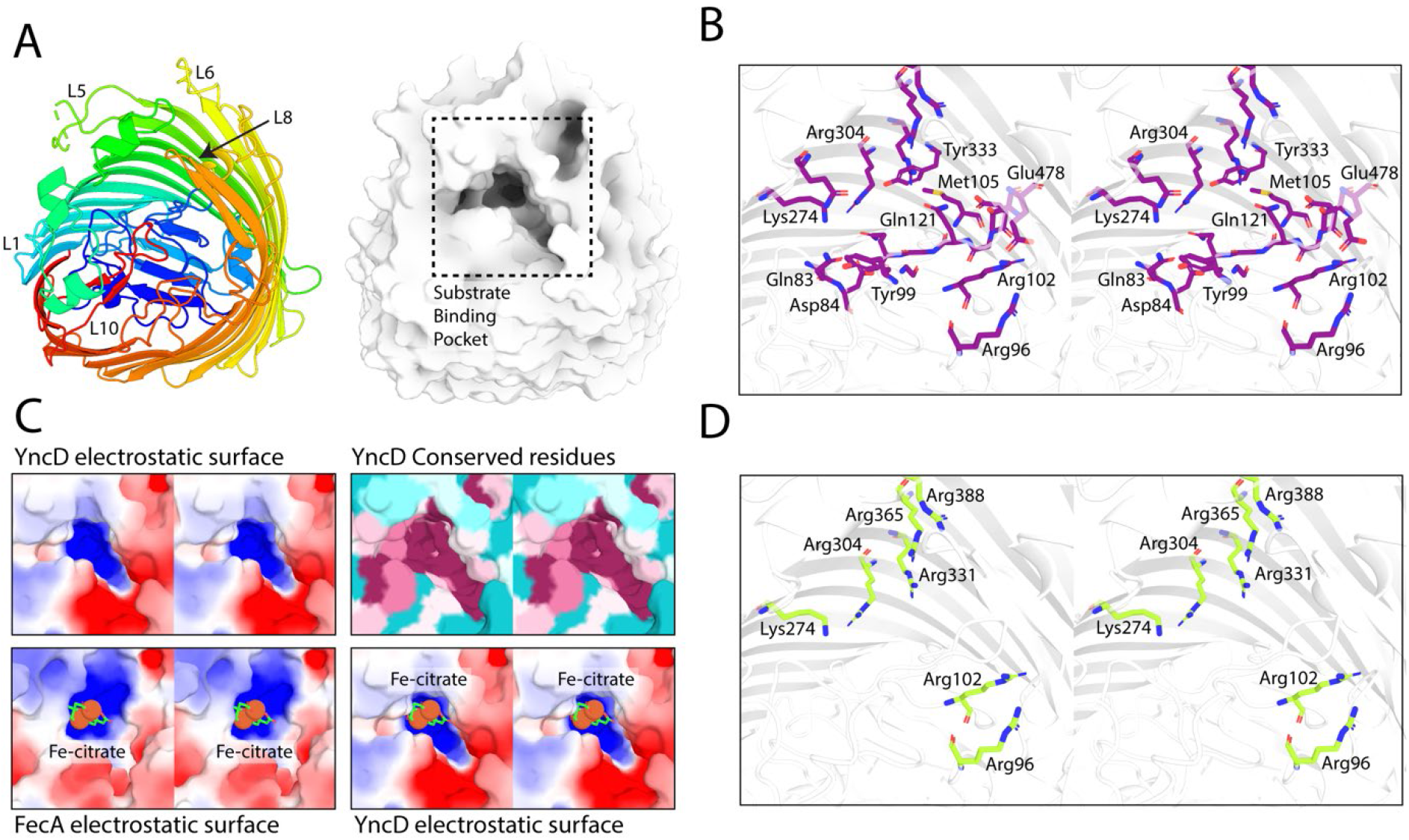
YncD possesses a conserved positively charged substrate-binding pocket. (A) A cartoon representation of YncD colored as in Figure 1A (left) and a surface representation in the same orientation showing the substrate-binding pocket (right). (B) A stereoview of the amino acids in the YncD substrate-binding pocket which are conserved in YncD homologues according to the program Consurf (33). The region shown corresponds to the boxed region in panel A, YncD is shown as a cartoon representation and conserved residues are shown as sticks. (C) A stereoview of the region of YncD from panel B shown as an electrostatic surface (top left), as surface colored according to Consurf conservation scores: dark purple = highly conserved, lighter purple = somewhat conserved, light blue = less conserved, dark blue = poorly conserved (top right). An electron static surface of FecA in complex with Fe-citrate shown as a stick and sphere representation (bottom left) and the position of Fe-citrate in the YncD binding pocket when the FecA-Fe-citrate complex with superimposed with YncD (bottom right). (D) A stereoview of positively charged conserved residues in the YncD substrate-binding pocket.

### YncD does not import ferric citrate or support growth of *E. coli* during iron limitation in pure culture

To test the hypothesis that YncD could support growth under iron-limiting conditions and specifically that it imports ferric citrate, we monitored the growth of an isogenic *E. coli* strain with and without YncD. Because TBDT iron-uptake systems are highly redundant, the *yncD* gene was deleted in a strain which is deficient in the 7 TBDTs involved in iron-uptake (Figure S4) (7). This ΔTBDT strain grows poorly under iron-limiting conditions (6), making any additional iron-acquisition defect easier to discern. In M9 minimal media, the growth of the ΔTBDT +/− *yncD* was identical, demonstrating that YncD does not play a role in iron acquisition under these conditions (Figure 4A). Since there is some residual free iron in M9 minimal media, the available iron was chelated with the addition of citrate or the monomeric catecholate compound 2,3-dihydroxybenzoic acid (DHB). As the ΔTBDT strain lacks the outer membrane transporters required to import these substrates, they limit the exogenous iron available to the bacteria. Accordingly, the growth of *E. coli* ΔTBDT was somewhat reduced in the presence of DHB and drastically reduced in the presence of citrate (Figure 4B). Deletion of *yncD* in this strain had no further effect on growth, demonstrating that YncD does not play a role in importing Fe-citrate or Fe-DHB under these conditions (Figure 4B). As expected growth of wildtype *E. coli* BW25113, which possesses TBDTs for both ferric-citrate (FecA) and Fe-DHB (Fiu, CirA, FepA) import (7,26) was identical in the presence or absence of these compounds (Figure 4C).

**Figure 4.**
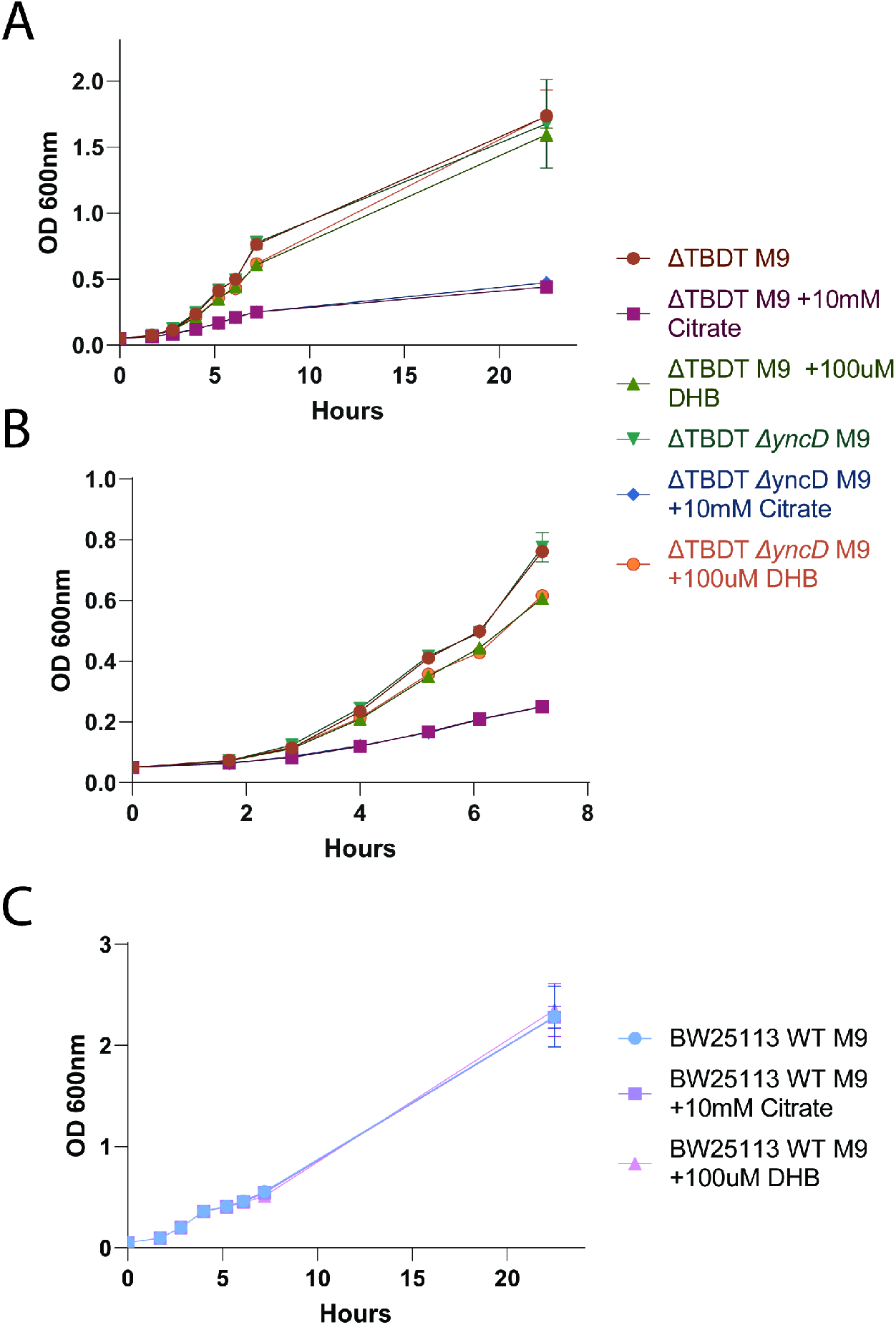
YncD does not import ferric citrate or assist in iron acquisition under laboratory conditions. (A) A growth curve of E*. coli* ΔTBDT and E*. coli* ΔTBDT/*ΔyncD* grown in M9 minimal media in the presence or absence of 10 mM citrate or 100 μM 2,3-dihydroxybenzoic acid. (B) A zoomed view of the growth curve from panel A hours 1 to 7. (C) the growth of wildtype *E. coli* BW25113 under the same conditions as the growth curve shown in panel A.

## Discussion

Determination of the function of proteins is one of the most challenging aspects of modern protein biochemistry. The expansion of genomic and metagenomic sequencing data has provided a wealth of proteins sequences, most of which are only partially characterised or of completely unknown function. Despite its difficulty, determining the function of novel proteins is invaluable, as without a robust understanding of the functional capacity of an organism’s proteome it is difficult to accurately predict its metabolic capabilities, physiology or lifestyle.

In this work, we show that YncD is a TBDT which is only distantly related to previously characterised members of this family and that YncD is widespread in gamma- and beta-proteobacteria. The crystal structure of YncD shows common structural features with the ferric-citrate transporter FecA, but from growth assays, YncD appears not to transport ferric-citrate. The structure of YncD represents the final structure of the 9 TBDTs possessed by the model bacterium *E. coli* K12 (6,7,23,26,34–36). This structure improves our understanding of the structural variation and similarities that exist in the TBDT family and paves the way for future studies addressing the function of YncD.

Seven of the nine TBDTs present in *E. coli* are involved in iron acquisition. All of these transporters are regulated by the Ferric uptake regulator (Fur) and are induced under iron-limiting conditions (37). BtuB, which imports the non-iron containing substrate cobalamin, is neither Fur regulated or induced under iron limiting conditions (37,38). Interestingly, like BtuB, YncD is neither Fur regulated or upregulated under iron-limiting conditions, suggesting that it may not function in the import of an iron-containing substrate (37,38). While the specific substrate transported by YncD remains to be determined, this work provides a number of clues and avenues for further research towards its identification. Based on its compact negatively-charged extracellular binding site, the substrate of YncD is likely to be relatively small and negatively charged.

As YncD is prevalent in bacterial species from soil and aquatic communities, its substrate is likely produced by members of these ecological communities. In addition, the impact of YncD on virulence phenotypes in *Salmonella* Typhi suggest that it is also a molecule which is relevant in a human host-context (28). Based on these considerations and structural parameters in the ligand-binding site, possible substrates for YncD would include molecules containing negatively charged phosphate groups. Other members of the TBDT family have been previously been implicated in thiamine pyrophosphate import through comparative genomics (39). Alternatively, YncD may import an organic acid containing compound in complex with a non-ferrous metal ion (26).

The importance of YncD for virulence in *Salmonella* Typhi (28), makes understanding the function of this transporter an important question for medical microbiology. Understanding the substrate of YncD and the role of this compound in bacterial virulence will improve our fundamental understanding of infection by this bacterium and provide a potential target for therapeutic intervention.

## Methods

### YncD Homologue Identification and Analysis

To determine the evolutionary relationship between YncD and other TBDTs of known structure and/or function these sequences were classified by an all-against-all BLAST clustering algorithm, based on pairwise similarities. The resulting data set was visualized with CLANS with an E-value cut off of 1×10^−10^ (29).

To identify YncD sequences in available bacterial genomes, a HMMER search was performed on the Uniprot reference proteomes database using YncD as the search sequence (30,31). An E-value cut-off of 1×10^−50^ applied to hits. This search yielded a total of 568 YncD homologue sequences. These sequences were classified by an all-against-all BLAST clustering algorithm, based on pairwise similarities. The resulting data set was visualized with CLANS with an E value cut off of 1×10^−130^. Sequence clusters were identified in CLANS using a network-based algorithm, with a minimum group size of 10 (29). The largest sequence cluster from this clustering contained 359 sequences, including YncD from *E. coli* and designated the YncD group. This group was isolated and subjected to further clustering with an E value cut off of 0. A multiple sequence alignment was performed on 331 full-length sequences from the YncD group using the clustal algorithm (40). Environment or host of isolation for a subset of YncD sequences (105 sequences) was determined from genome metadata from the Ensemble and Uniprot databases (41,42).

### YncD Expression and Purification

The open reading frame encoding YncD was amplified by PCR from *E. coli* BW25113 using primers containing 5’ BamHI and 3’ XhoI restriction sites (Forward: CCATCGGATCCGGCTGATGAACAGACTATGATTGTC, Reverse: CCATCCTCGAGTTACTCAAATCTCCACGCAATATTCAT). This YncD open reading frame was then cloned into a modified pET20b vector with a PelB signal sequence, followed by an N-terminal 10 × His-tag and TEV cleavage site, via restriction digestion and ligation. The resulting vector was transformed into *E. coli* BL21 (DE3) C41 cells (43). Protein expression was performed in terrific broth (12 g tryptone, 24 g yeast extract, 61.3 g K_2_HPO_4_, 11.55 g KH_2_PO_4_, 10 g glycerol) with 100 mg.ml^−1^ ampicillin for selection. Cells were grown at 37 °C until OD_600_ of 1.0, induced with 0.3 mM IPTG, and grown for a further 14 hours at 25 °C. Cells were harvested by centrifugation and lysed using a cell disruptor (Emulseflex) in Lysis Buffer (50 mM Tris, 200 mM NaCl [pH 7.9]) in the presence of 0.1 mg.ml^−1^ Lysozyme, 0.05 mg.ml^−1^ DNAse1 and complete protease cocktail inhibitor tablets (Roche).

The resulting lysate was clarified by centrifugation at 10,000 g for 10 min, the supernatant from this low-speed spin was then centrifuged to 1 h at 100,000 g to isolate a membrane fraction. The supernatant was decanted, and the membrane pellet was suspended in lysis buffer using a tight-fitting homogeniser. The resuspended membranes were solubilised by the addition of 10 % Elugent (Santa Cruz Biotechnology) and incubated with gentle stirring at room temperature for 20 min. The solubilised membrane protein fraction was clarified by centrifugation at 20,000 g for 10 min. The supernatant containing the solubilized proteins was applied to Ni-agarose resin equilibrated in Ni-binding buffer DDM (50 mM Tris, 500 mM NaCl, 20 mM Imidazole, 0.03% Dodecylmaltoside (DDM) [pH7.9]). The resin was washed with 10-20 column volumes of Ni binding buffer DDM before elution of the protein with a step gradient of, 10, 25 and 50, 100 % Ni gradient buffer DDM (50 mM Tris, 500 mM NaCl, 1 M Imidazole, 0.03 % DDM [pH7.9]). YncD eluted at the 50 and 100 % gradient steps. Eluted fractions containing YncD were pooled and applied to a 26/600 S200 Superdex size exclusion column equilibrated in SEC buffer DDM (50 mM Tris, 200 mM NaCl, 0.03 % DDM [pH 7.9]). To exchange YncD into the detergent Octyl β-D-glucopyranoside (βOG) for crystallographic and biochemical analysis, SEC fractions containing YncD were pooled and applied to Ni-agarose resin, equilibrated in βOG buffer (50 mM Tris, 200 mM NaCl, 0.8 % βOG [pH 7.9]). The resin was washed with 10 column volumes of βOG buffer before elution with βOG buffer + 250 mM imidazole. Fractions containing YncD were pooled, and 6 × His-tagged TEV protease (final concentration 2 mg.ml^−1^) and DTT (final concentration 1 mM) were added. This solution was then dialysed against of βOG buffer at 4-6h at 20 °C to allow TEV cleavage of the 10 × his-tag from YncD and removal of excess imidazole. The sample was then applied to Ni-agarose resin to remove TEV protease and the cleaved histidine-containing peptide. The flow-through containing YncD from this step was collected, concentrated to 10 mg.ml^−1^ in a 30 kDa cut-off centrifugal concentrator, snap-frozen and stored at −80 °C.

### YncD Crystallisation, Data Collection and Structure Solution

Purified YncD in βOG was screened for crystallisation conditions using commercially available screens (approximately 600 conditions). Crystals grew in a number of conditions, with a condition containing 0.2 M Ca acetate, 0.10 M Tris pH 8.5 and 20 % PEG 3000 chosen for optimisation. Concentration and pH grid screening was performed yielding an optimised crystallisation conditions containing 0.17 M Ca acetate, 0.10 M Tris pH 7.0 and 16 % PEG 3000. Crystals from this condition were looped, crystallisation solution was removed by wicking and crystals were flash-cooled in liquid N_2_. Diffraction data were collected at 100 K at the Australian synchrotron and processed in the space group P2_1_22_1_ to 3.2 Å. Molecular replacement of this dataset using the structures of various TBDTs as a starting model failed and so soaking was performed to obtain a heavy atom derivative. Using a fine acupuncture needle a small quantity of potassium tetranitroplatinate salt was added to the drop containing YncD crystals, the well was resealed and crystal incubated for 60 minutes. These crystals were then were looped, crystallisation solution was removed by wicking and crystals were flash-frozen in liquid N_2_. Data was collected from the potassium tetranitroplatinate soaked crystals at 100 K at the Australian synchrotron using a wavelength of 0.987 Å, with crystals diffracting to 3.5 Å. A heavy atom search was performed by SAD using Shelx within the CCP4 software package (44,45). Three heavy atom sites were identified, which were provided to Autosol withing the Phenix software package for phasing and density modification (46). An initial structure of YncD was constructed using these experimentally-phased density modified maps using Coot (47). This model was used for molecular replacement of the 3.2 Å native data using phaser (48). The model of YncD was then build and refined using Coot, Phenix package and Buster (46,47,49).

### *E. coli ΔyncD* Mutant Generation and Growth Analysis

The *E. coli* BW25113 *ΔyncD* mutant strain was created using the λ Red system (50). A Kanamycin-resistance cassette flanked by 300 bp of genomic DNA either side of the chromosomal location of *yncD* was amplified by PCR (primers: F: AAACAGGCTATTTCGCTTAGCGA, R: GAACCTAACAGTAATGAACCACG using the specific mutant from the *E. coli* Keio collection (51) as a template, generating the yncD-Kan KO cassette. *E. coli* ΔTBDT (6) was transformed with the λ Red recombinase plasmid pKD46 (50) and grown at 30 °C (LB broth + 100 μg.ml^−1^ ampicillin + 100 uM Fe(II)SO_4_) to an OD600 nm of 0.1 before λ Red recombinase was induced by the addition of 0.2% L-arabinose. The culture was then grown at 30 °C until an OD600 nm of 0.6–0.8 was attained and were transformed with the yncD-Kan KO cassette using the room-temperature electroporation method (52). Briefly, bacterial cells were isolated by centrifugation at 3000 g for 3 min and washed twice with a volume of sterile 10% glycerol equal to the volume of culture used. The cells were then resuspended in 10% glycerol to a volume of 1/15 of that of the culture. The yncD-Kan KO cassette DNA (100–500 ng) was then added to 100 μl of the resuspended bacteria and the mixture was electroporated. 1 ml of LB broth was added to the cells post-electroporation, and the culture was recovered at 37°C for 1 h before plating onto LB agar + 30 μg ml^−1^ kanamycin + 100 uM Fe(II)SO_4_. PCR was used to validate that colonies did indeed have the KanR cassette in place of the gene of interest.

To remove the KanR gene and generate a “clean” *yncD* deletion, the mutant strain was transformed with the plasmid pCP20 (53) containing the ‘flippase cassette’. Cells were grown at 30 °C under either ampicillin (100 μg ml^−1^) or chloramphenicol (30 μg ml^−1^) selection to maintain the plasmid. A single colony of the mutant pCP20-containing strain was used to inoculate 1 ml LB broth + 100 uM Fe(II)SO_4_ (no selection). The culture was grown overnight at 43°C to activate expression of the flippase gene. This culture was then subjected to tenfold serial dilution in sterile LB and plated onto LB agar + 100 uM Fe(II)SO_4_ with no selection. The resulting colonies were patched onto LB agar containing kanamycin, chloramphenicol or no selection. PCR was used to validate colonies that, while unable to grow in the presence of kanamycin or chloramphenicol, grew in the absence of selection and had no remnant of the KanR cassette in the deletion of *yncD*. *E. coli* ΔTBDT, *E. coli* ΔTBDT/*ΔyncD* and wildtype *E. coli* BW25113 were grown in LB broth until stationary phase. These cultures were used to inoculate 20 ml of M9 minimal media +/− 10 mM citrate or +/− 100 μM 2,3-Dihydrobenzoic acid, to an OD600nm of 0.05. Cultures were grown at 37 °C with shaking and the rate of growth was quantified by measuring OD600nm at hourly intervals.

## Data availability

The crystallographic coordinates and associated structure factors of YncD produced in this study are available in the Protein Data Bank (PDB) with the accession code: 6V81

## Acknowledgments

This research was undertaken in part using the MX2 beamline at the Australian Synchrotron, part of ANSTO, and made use of the Australian Cancer Research Foundation (ACRF) detector. This research was undertaken on the MX2 beamline at the Australian Synchrotron, part of ANSTO (CAP12312). We would like to thank the Monash Crystallisation Facility for their assistance with sample characterization, crystallographic screening, and optimization. The work was funded by the Australian Research Council (ARC; FL130100038) and the National Health & Medical Research Council (NHMRC Program in Cellular Microbiology, 1092262). R.G. was funded by a Sir Henry Wellcome Fellowship award (106077/Z/14/Z). T.L. is an ARC Australian Laureate Fellow (FL130100038).

## Author contributions

Conceived and designed the experiments: RG, TL

Performed the experiments: RG

Analyzed the data: RG

Contributed reagents/materials/analysis tools: RG, TL

Wrote the paper: RG

