## Supplemental Data for "The crystal structure of the TonB-dependent transporter YncD reveals a positively charged substrate binding site"

**Data S1: A multiple sequence alignment of YncD sequences from clustering analysis outlined in Table S1 and 2**

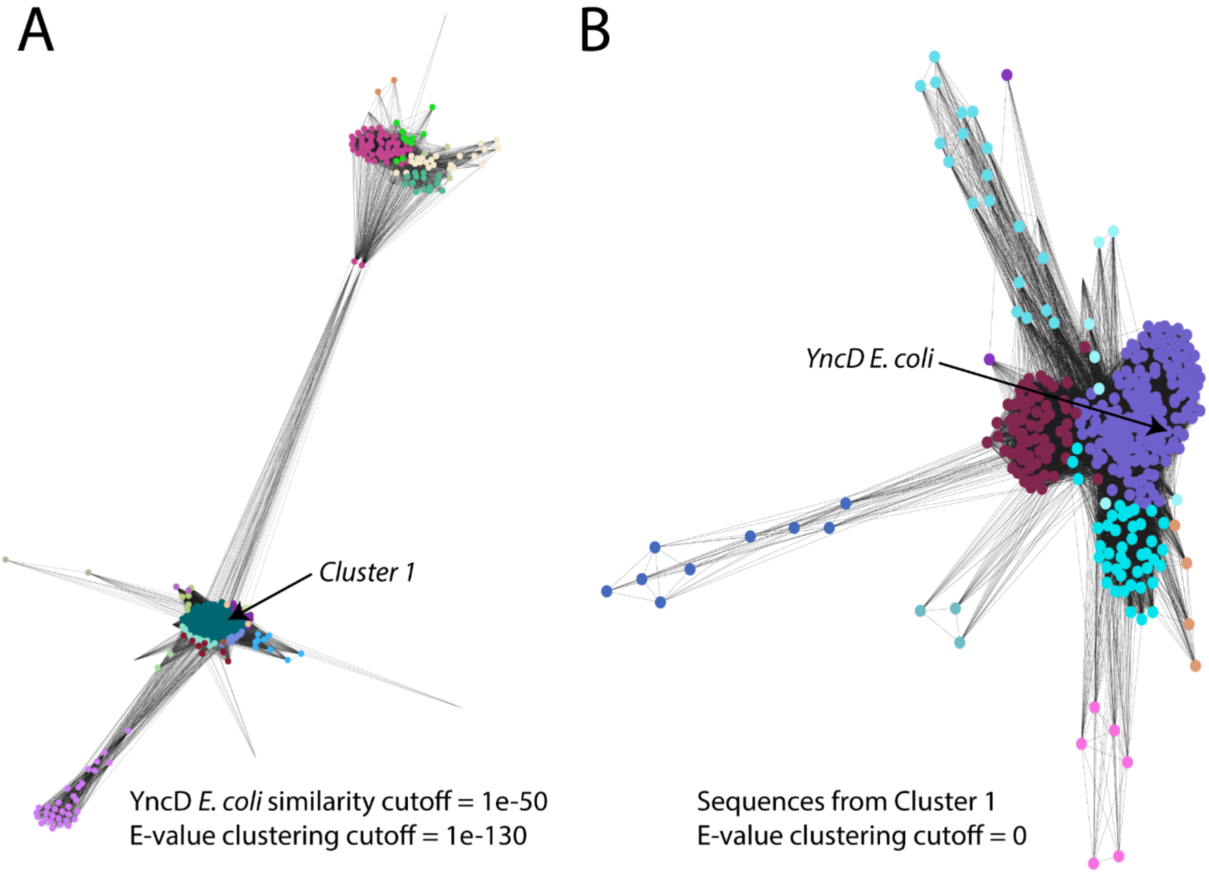


**Figure S1 CLANS sequence clustering of YncD homologues from diverse bacterial isolates.** (A) CLANS clustering of YncD homologues from the Uniprot references proteomes database identified with an E-value cutoff of $1x10^{-50}$ and clustered based on E-value cutoff of $1x10^{-130}$ or smaller. (B) YncD sequences from cluster 1 in panel A clustered based on E-value cutoff of 0.

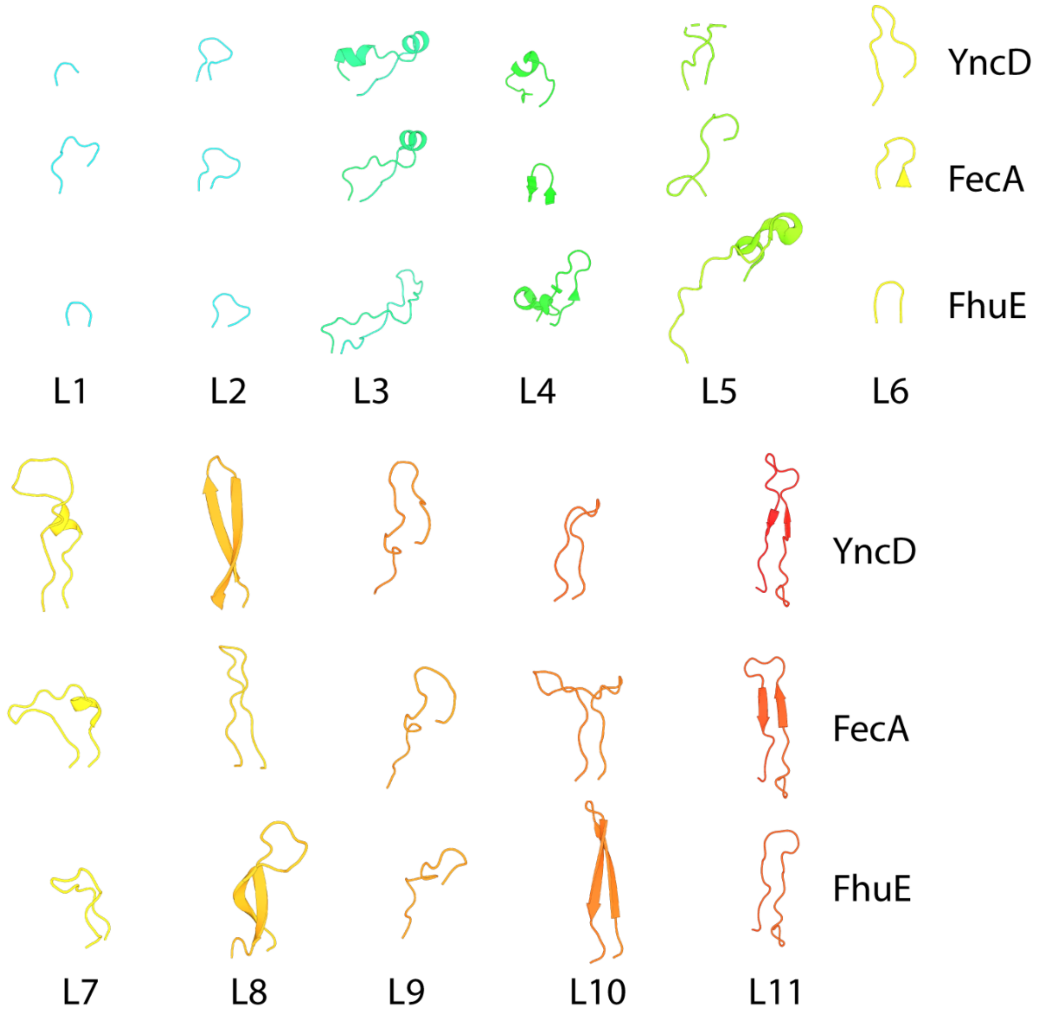


**Figure S2 Comparison of the extracellular loops of YncD with other TBDTs.** A dissection of the extracellular loops of YncD and the TBDTs FecA and FhuE, providing an indication of the similarity of secondary structure between the extracellular loops of these transporters.

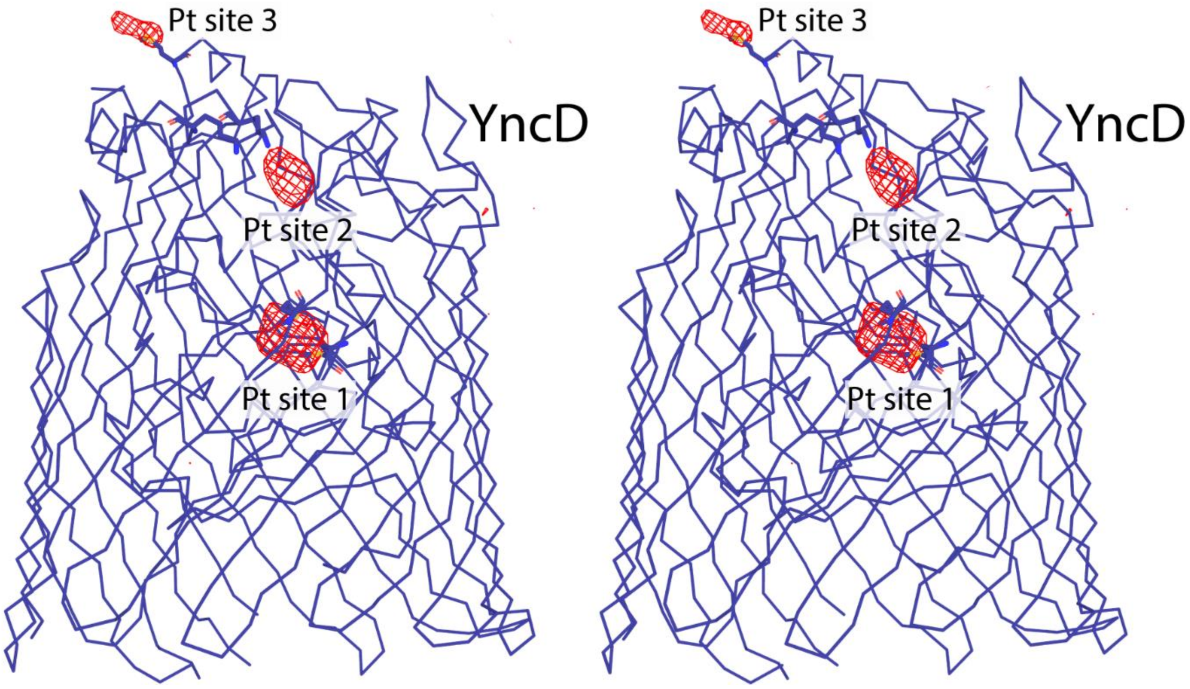


**Figure S3 A stereoview of the anomalous difference map derived from diffraction data collected from a potassium tetranitroplatinate soaked YncD crystal.** Peaks in the anomalous difference map showing binding sites of tetranitroplatinate molecules are shown as red mesh contoured to 4 σ. Pt sites 1 and 2 represent high occupancy sites > 0.5, and Pt site 3 represents a low occupancy site < 0.2. YncD is shown as a blue ribbon representation.

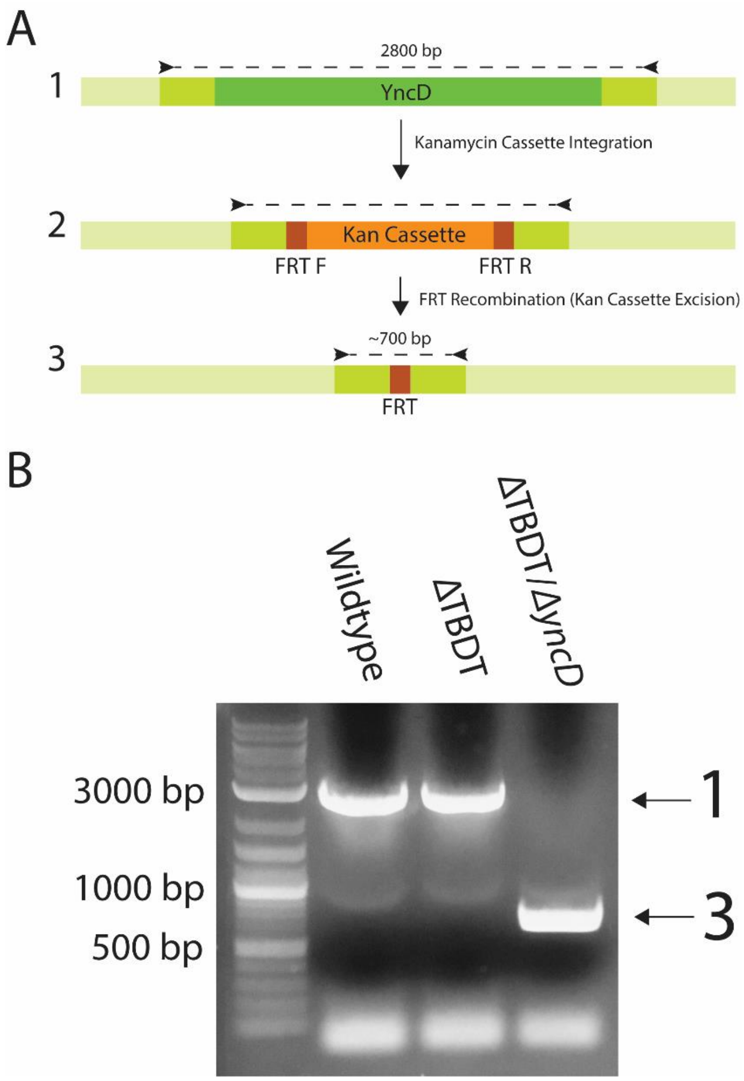


**Figure S4 Creation of the *E. coli* ΔTBDT/*ΔyncD* strain.** (A) A schematic of the *yncD* knockout process, showing step 1 the original *yncD* gene, step 2 the replacement of the *yncD* gene with a kanamycin resistance cassette (Kan cassette) and step 3 removal of Kan cassette via FRT-site mediated recombination. The primers utilised for Kan cassette amplication and for confirmation of knock out creation are shown as black arrows. (B) PCR amplification from genomic DNA of wildtype, ΔTBDT, and ΔTBDT/*ΔyncD E. coli* strains using *yncD* KO primers shown in panel A, with amplification product corresponding to sizes predicted in the KO scheme. The genome of the ΔTBDT/*ΔyncD* strain was sequenced confirming no non-desired mutations were introduced.

**Table S1 YncD sequences from diverse organisms identified by HMMER search and clustering analysis** (Attached File)

**Table S2 Sequence identity matrix of full length YncD sequences outlined in Table S1** (Attached File)

**Table S3 YncD structural homologues identified by the Dali server [1]**

| Chain | Z | rmsd | lali | nres | %id | Description |
|---|---|---|---|---|---|---|
| 1kmo-A | 36.6 | 2.2 | 596 | 661 | 21 | Iron(iii) dicitrate transport protein FecA |
| 2gsk-A | 36.3 | 2.7 | 531 | 590 | 18 | Vitamin B12 transporter BtuB |
| 4epa-A | 34.3 | 2.7 | 589 | 632 | 20 | Ferric-yersiniabactin transporter FyuA |
| 2o5p-A | 32.3 | 2.8 | 577 | 772 | 18 | Ferripyoverdine Receptor |
| 1fi1-A | 32.2 | 2.4 | 589 | 707 | 20 | Ferrichrome-Iron Receptor |
| 5m9b-A | 30.9 | 3 | 543 | 707 | 19 | Ferric-Enterobactin Receptor |
| 5fq6-M | 30.8 | 3.6 | 573 | 948 | 16 | SusC |
| 6h7f-A | 30.7 | 2.8 | 582 | 674 | 15 | BauA |
| 6fok-A | 29.7 | 3.1 | 571 | 651 | 18 | Copper Transporter OprC |
| 6bpm-A | 29.6 | 2.8 | 581 | 711 | 20 | Catecholate Siderophore Receptor Fiu |
| 3csl-A | 29.2 | 2.8 | 568 | 753 | 18 | HasR |
| 4aip-B | 28.4 | 3.1 | 557 | 665 | 17 | FrpB |
| 3v89-A | 27.9 | 3.4 | 567 | 853 | 16 | TbpA |
| 5fr8-A | 27.8 | 2.8 | 542 | 707 | 20 | Catecholate Siderophore Receptor PirA |
| 4rdr-A | 27.7 | 3.1 | 556 | 706 | 15 | ZnuD |
| 4zgv-A | 27.2 | 3.7 | 558 | 809 | 14 | Ferredoxin Transporter FusA |
| 6ofr-A | 25.5 | 3.5 | 556 | 758 | 16 | Putaative Protein Transporter YddB |
